# Rapid identification of African swine fever virus in diagnostic samples using CRISPR-Cas

**DOI:** 10.1101/2024.12.27.630508

**Authors:** Sekhar Kambakam, Julia Thomas, Suelee Robbe-Austerman, Karthik Shanmuganatham, Rachel Palinski

## Abstract

African Swine Fever Virus (ASFV) is a high consequence, highly transmissible pathogen affecting swine causing African Swine Fever (ASF), a devastating disease, with high mortality rates in naive populations. Due to the likelihood of significant economic impacts associated with an ASF outbreak, considerable resources have been allocated in the United States (U.S.) to safeguard the swine industry against this threat. Ongoing outbreaks of ASF in the Dominican Republic and Haiti further threaten U.S. swine due to their proximity and involvement in movement to and from North America. While surveillance programs are ongoing, there are limited point-of-care (POC) tests available during outbreaks that maintain the sensitivity and specificity standards of laboratory testing (*e.g*., qPCR). However, the recently developed CRISPR-Cas testing systems may provide comparable high-quality results. In a CRISPR-based diagnostic assay, CRISPR effectors can be programmed with CRISPR-RNA (crRNA) to target specific DNA or RNA. Upon target binding, the Cas enzyme undergoes collateral cleavage of nearby fluorescently quenched reporter molecules (ssDNA or ssRNA), which can be detected under blue light or a fluorescence microplate reader. Furthermore, this tool is rapid, simple, cost-effective and can be performed with inexpensive equipment. For these reasons, we sought to develop a low-cost visual detection method for ASFV by employing the recombinase polymerase amplification (RPA)-dependent CRISPR-Cas12a technique that can be utilized in the field as a point-of-care-assay. Our CRISPR-Cas12a assay demonstrated comparable sensitivity and specificity to qPCR, both visually and when quantified using a fluorescent reader. In whole blood samples from ASFV-suspect or ASFV-negative cases, the CRISPR assay achieved a sensitivity of 98.3% (10^2^ DNA copies) and a specificity of 100%. Finally, an assessment of the reaction time constraints indicated that results can be visualized in as little as seven minutes with a peak fluorescence at 40 min (RPA and CRISPR steps). The results of this feasibility assay validation allow for the rapid development of sensitive and specific POC tests that may be used for outbreak response in the future.

## INTRODUCTION

African Swine Fever Virus (ASFV) possesses a complex architecture, composed of 180-190 kilobase pairs (kb) of double-stranded DNA. The genome encodes over 160 open reading frames (ORFs) and belongs to the genus *Asfivirus* and is the sole member of the *Asfarviridae* family (Chapman et al., 2011; Alonso et al., 2018; Bohorquez et al., 2023; Borca et al., 2024). ASFV is highly contagious in domestic and wild swine species (Pollock et al., 2021; Li et al., 2022). ASFV can be transmitted by direct contact but is stable in the environment, feedstuffs, equipment, and fomites for an extended period of time. ASF cause a wide range of clinical signs including fever, lethargy, loss of appetite, vomiting, diarrhea, hemorrhage, and respiratory distress (Li et al., 2022; Wu P et al., 2024). Clinical disease in naïve animals is often severe, with a mortality rate up to 100% in infected animals. Once infected with highly virulent ASFV isolates, pigs can die within 7–10 days after the onset of symptoms (Salguero, 2020).

African Swine Fever (ASF) is believed to have originated in sub-Saharan Africa where it remains endemic, circulating in both wild and domestic pigs and causes sporadic outbreaks. Historically, ASF outbreaks have occurred outside of Africa, notably in Europe and the Americas; and most of these outbreaks were eradicated. In 2007, an ASF outbreak in Georgia led to the virus becoming endemic in the wild boar populations in Eastern Europe (http://www.promedmail.org, archive no. 20070607.1845; Rowlands et al., 2008). For over 10 years, the virus remained undetected outside of these regions; however, in 2018, a breach in biosecurity practices led to ASF being introduced into the commercial naive swine population in China. The virus spread rapidly, causing an economic collapse of the swine industry in the country (Zhao et al., 2019). Shortly thereafter, in 2021, ASFV was detected on the Caribbean Island of Hispaniola. (Schambow et al., 2022). This outbreak remains ongoing (Schambow et al., 2022; Bohorquez et al., 2023; Borca et al., 2024; World Organisation for Animal Health -WOAH, ASF-situation reports 57, 2024), posing a significant threat to the United States (U.S.) swine industry. For these reasons, the U.S. significantly increased the ASF surveillance program within the continental U.S. developed a protection zone around the U.S. territories of Puerto Rico and the U.S. Virgin Islands that halted the movement of pork and pork products from the territories into the continental U.S. assisted in adding ASF testing in a veterinary laboratory in Puerto Rico, and has been working with the Dominican Republic in implementing control measures. Islands in the Caribbean are challenged in that sophisticated veterinary laboratory resources are limited, and transportation of samples to a main laboratory can be challenging. For these reasons, it is crucial to develop and deploy rapid, simple, point-of-care (POC) ASF detection assays that are comparable to gold standard laboratory diagnostic tests to aid in outbreak control and prevention efforts.

To date, qPCR targeting the highly conserved *p72* viral gene has been the gold-standard molecular diagnostic assay for the detection of ASFV. While qPCR is sensitive and specific, it requires expensive equipment, reagents, trained personnel, and often specialized infrastructure to perform the assay. Other techniques, such as recombinase polymerase amplification (RPA) and loop-mediated isothermal amplification (LAMP), have been tested as alternatives to qPCR in the field (Miao et al., 2019; Zhai et al., 2020, James et al., 2010; Mee et al., 2020; Bohorquez et al., 2023); but these techniques have sensitivities and specificities below that of the gold-standard qPCR (Kim et al., 2023). The introduction of Clustered Regularly Interspaced Short Palindromic Repeats (CRISPR) and CRISPR-associated protein (CRISPR-Cas) technology has revolutionized the field of genetic engineering and has potential applications in various areas such as gene therapy, gene activation, gene silencing, drug delivery, and more (Li et al., 2023). CRISPR-Cas nuclease cleavage activity can be leveraged as a next-generation diagnostic assay for *in vitro* nucleic acid detection. When combined with isothermal amplification, it produces exceptional point-of-care diagnostic tests (Ghouneimy et al., 2023; Gootenberg et al., 2017, 2018; Chen et al., 2018).

CRISPR-Cas-based tests have been assessed for a wide range of pathogens, such as ASFV (Ren et al., 2021; Fu et al. 2021; Qin et al., 2022; Qian et al., 2022), porcine reproductive and respiratory syndrome virus (PRRSV; Liu et al., 2021), tobacco curly shoot virus (TCSV; Smith et al., 2020), *Listeria monocytogenes* (Li et al., 2021), foodborne bacteria (*Escherichia coli* and *Streptococcus aureus*; Wang et al., 2020), *Salmonella* spp. (An et al., 2021), *Mycobacterium tuberculosis* (Ai et al., 2019), *Phytophthora sojae* (Guo et al., 2023), and shrimp viruses (Major et al., 2023), the results of which are consistent with qPCR standards. More importantly, during the COVID pandemic, the FDA approved the use of CRISPR-based diagnostic assays for SARS-CoV-2 detection allowing CRISPR to gain a foothold in the commercial POC industry and providing an opportunity to develop similar assays against other pathogens (Yoshimi et al., 2020; Joung et al., 2020; Kellner et al., 2019).

In this study, we developed a distinct ASFV-RPA-CRISPR assay by selecting and testing the designed crRNA (CRISPR-RNA) effectiveness and crRNA-Cas12a complex cleavage specificity. We then optimized the assay to assess the sensitivity and specificity and verified its effectiveness across different biological matrices compared to the standard ASFV qPCR assay. Furthermore, we demonstrated diagnostic sensitivity and specificity in clinical samples. The results of this study provide the groundwork for a large-scale POC assay assessment for ASFV detection as well as a robust workflow that works with commonly submitted diagnostic samples.

## MATERIALS AND METHODS

### ASFV-RPA primers and CRISPR reagents

RPA primers were designed according to the TwistAmp Assay design manual (TwistDx Limited, Maidenhead, UK) and obtained from IDT (Integrated DNA Technologies Inc., Coralville, IA, U.S., Table 1). We designed and synthesized two crRNAs from IDT to compare the efficiency of the RPA-CRISPR-Cas12a assay with the qPCR results. Primers and crRNA sequences targeting the ASFV-*p72* gene are listed in Table 1. The reporter used in this design was a 5’-56-FAM and the quencher was 3’-BHQ.

**TABLE 1.**
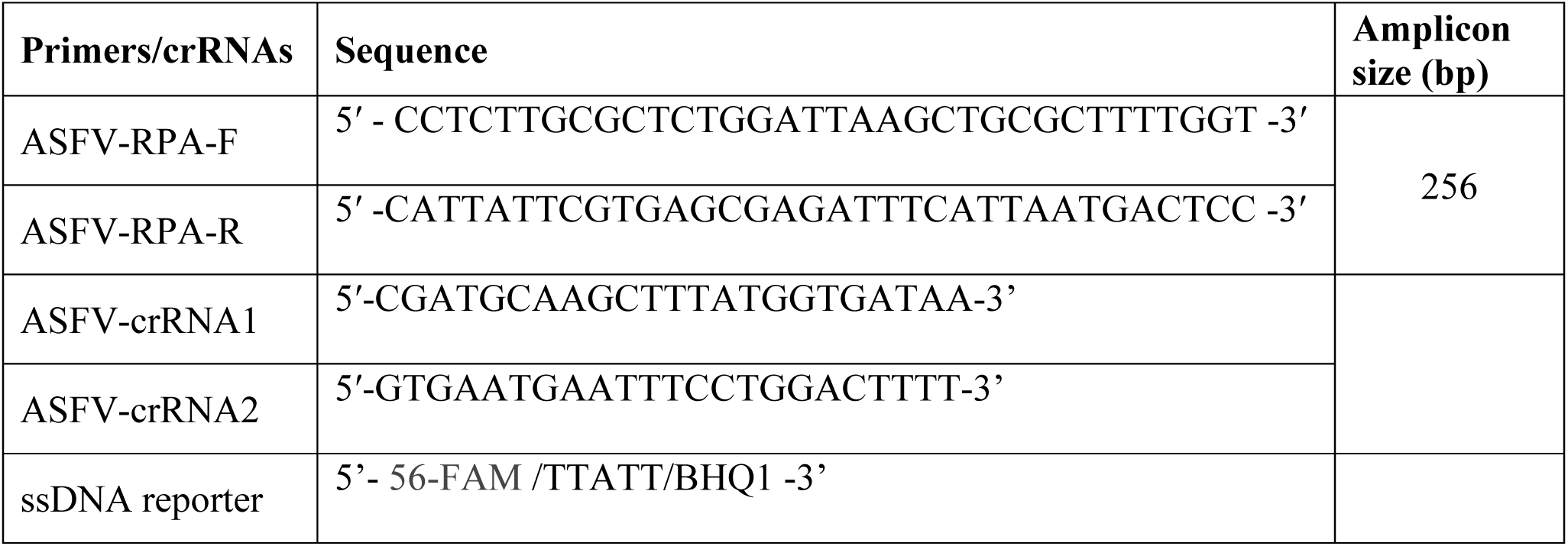
RPA primers, crRNAs, and reporter used in this study.

### Controls and nucleic acids for specificity assessment

In this study, we selected a conserved region of the *p72* gene (Genbank # MN886930; 621 to 1296 bp) for synthesis and cloning into a pUCIDT plasmid vector obtained from IDT (Coralville, IA, U.S.). The recombinant *p72* plasmid DNA is termed *p-p72* and is used as a positive control throughout the study. For the assessment of specificity, DNA or cDNA was obtained from the USDA-Center for Veterinary Biologics then quantities and qualities assessed by Nanodrop before use in this optimization. Nucleic acid obtained included swine pox virus (SwPV), modified vaccinia ankara virus (MVAV), porcine epidemic diarrhea virus (PEDV), porcine hemagglutinating encephalomyelitis virus (PHEV), bovine viral diarrhea virus (BVDV), and porcine reproductive and respiratory syndrome virus (PRRSV). Double distilled water was utilized as a negative control throughout these experiments.

### RPA-dependent CRISPR-Cas12a assay setup

The RPA-CRISPR-Cas12a ASFV assay was designed as a single-tube, two-step reaction using a TwistAmp (TwistDx Limited, Maidenhead, UK) pellet. First, TwistAmp pellets, 480 nM of ASFV RPA F and ASFV RPA R primers, 29.5 uL of the TwistAmp rehydration buffer, 14 nM magnesium acetate, and 7.2 uL of water were added to a tube and pipette-mixed on ice. After mixing, the tube was spun down briefly, and 10 uL of the mixture was aliquoted into PCR tubes. Five uL of either template or control (water or plasmid DNA) were added to the PCR tubes. The PCR tubes were mixed and incubated at 37°C for 2-30 minutes (2, 5, 10, 15, 20, 25, and 30 minutes). After incubation, 50 nM of Cas12a (LbaCas12a, Cpf1 enzyme), 75 nM of crRNA, 0.5 uM of ssDNA reporter, and 1X NEB buffer 2.1 (NEB, Ipswich, MA, U.S.) were added to the RPA reaction tube. The RPA-CRISPR-Cas12a reaction tubes were incubated at 37°C for 1-30 minutes (1, 2, 5, 10, 15, 20, 25, and 30 minutes). Fluorescence was visualized using the Invitrogen E-gel dock blue light transilluminator (ThermoFisher Scientific, Waltham, MA, U.S.) and quantified on a Tecan Cytospark multimodal plate reader (Seestrasse, Switzerland) unless otherwise specified.

### Nucleic acids extraction and qPCR assay

For negative surveillance diagnostic samples, nucleic acids were extracted using the MagMAX CORE Nucleic Acid Purification Kit on a KingFisher Flex instrument as specified by the manufacturer (ThermoFisher Scientific, Waltham, MA, U.S.). For suspect blood samples, we manually extracted the nucleic acid using the MagMAX CORE Nucleic Acid Purification Kit in accordance with the manual instructions. ASFV qPCR was performed on the extracted nucleic acids as specified by the National Animal Health Laboratory Network standard protocol (Zsak et al., 2005). Ct values below 40 were considered positive. The Ct value of 45 was considered as no amplification.

### Analytical sensitivity and specificity

To assess analytical sensitivity, *p72* plasmid DNA was diluted 1:10 in sterile H_2_O ten times starting at 10^10^ copies. Analytical specificity was determined with 20 to 50 ng of nucleic acid from SwPV, MVAV, PEDV, BVDV or PRRSV. Matrix effects on the RPA-CRISPR-Cas12a ASFV assay were determined by spiking 10^6^ copies of *p72* plasmid DNA in either whole blood, serum, or a blood swab in MTM (Longhorn Vaccines & Diagnostics LLC, Bethesda, MD, U.S.).

### Diagnostic sensitivity and specificity

Negative surveillance diagnostic samples (39 total) were obtained from the continental US (CONUS) ASFV feral swine surveillance program as blood swabs in MTM. ASFV suspect whole blood samples (66 total; 47 positive and 19 negative) with an unconfirmed ASF status were obtained from the Laboratorio Veterinario Central (LAVECEN) in the Dominican Republic. These samples were collected as part of routine ASFV surveillance and depopulation efforts. We used qPCR to confirm the negative and positive results of both negative and suspect blood samples before testing the RPA-CRISPR-Cas12a assay. The samples were processed and tested at the biosafety level 3 (BSL-3) facility at the USDA National Centers for Animal Health (NCAH) in compliance with USDA and Department of Health and Human Services (HHS) safety regulations regarding select agents. ASFV positive samples were visualized using a MyGel InstaView gel dock (ThermoFisher Scientific, Waltham, MA, U.S.) and fluorescence was quantified by assessing photographs on ImageJ.

### Statistical analysis

The standard error of the mean (SEM) was calculated from three independent replicates. For statistical significance, unpaired two-tailed student’s t-tests were performed using GraphPad Prism 10 software. The confidence intervals for diagnostic sensitivity and specificity were calculated using the C.I. Calculator (https://www2.ccrb.cuhk.edu.hk/stat/confidence%20interval/Diagnostic%20Statistic.htm; Centre for Clinical Research and Biosatatistics).

## RESULTS

### Molecular detection of ASFV visually using an RPA-CRISPR-Cas12a-mediated tool

We evaluated a visual detection assay for ASFV using a single-tube, two-step reaction that combines both RPA and CRISPR techniques. The pre-mixed CRISPR reagents were employed on the RPA amplicons to visualize fluorescence resulting from crRNA-Cas12a-mediated collateral cleavage of the fluorescence reporter (Fig. 1A). We designed two independent crRNAs targeting the *p72* gene of ASFV and assessed their dependent Cas12a trans-cleavage efficiency to detect amplicons from the RPA reaction (Figs. 1B and 2). Both crRNA1 and crRNA2-Cas12a reactions displayed strong fluorescence signals, with the crRNA1 resulting in a significantly higher (∼12%) fluorescence signal than crRNA2. From these results, crRNA1 was utilized for further experiments (Fig. 2). In both experiments, the control did not display a visible fluorescence signal (Fig. 2).

**FIG 1.**
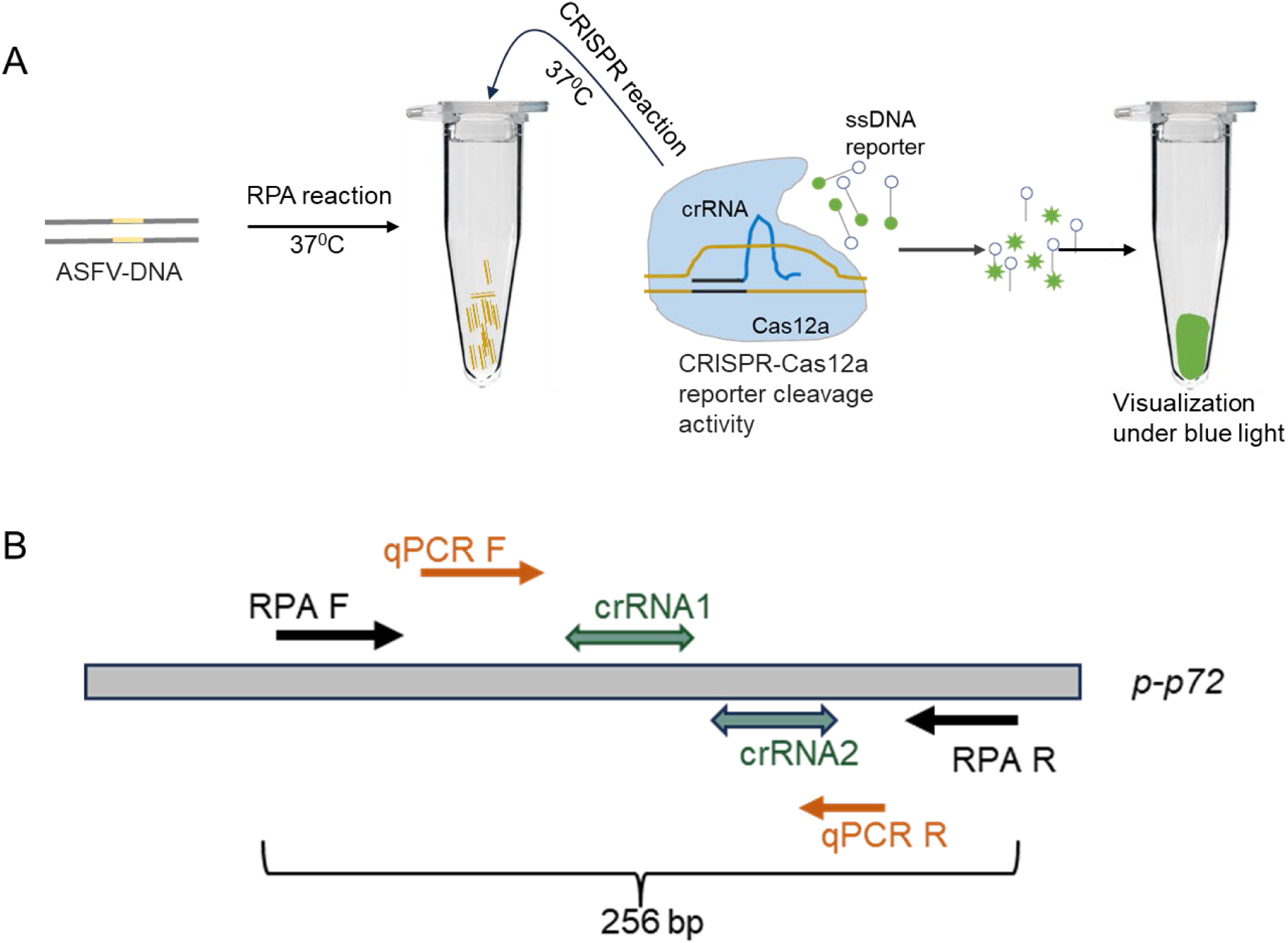
The RPA-dependent CRISPR-based diagnostic assay for the visual detection of ASFV. A. The target region is amplified in an RPA reaction at 37°C. The Cas12a cleaves the fluorescence reporter and quencher-labeled ssDNA molecule when combined with the target amplicon producing a visible fluorescence that can be seen using a common blue light transilluminator. This fluorescence is quantified on a Tecan Spark Multimode plate reader or in ImageJ. B. Schematic representation of RPA, PCR primers, and crRNA positions on the *p72* (B646L) gene fragment, which is cloned in the plasmid vector, *p-p72*. The RPA reaction yields an amplicon size of 256 bp.

**FIG 2.**
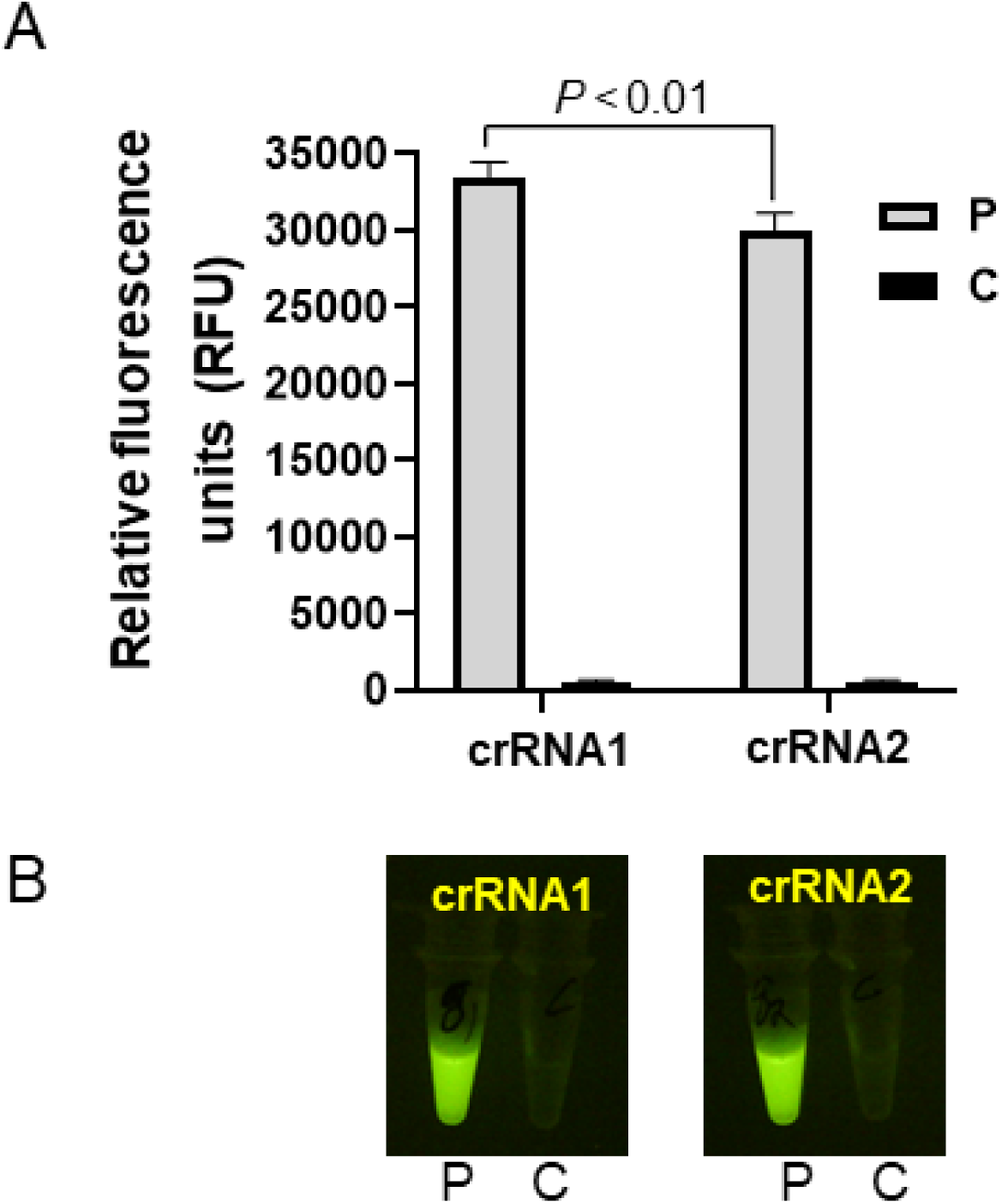
Validation of crRNA efficiency. A. The fluorescence signal was quantified using a Tecan Spark Multimode microplate reader. The data are mean ± SEM from the three replicates. P values were calculated using a Student’s t-test. B. The resulting fluorescence was observed using a blue light transilluminator. P represents *p-p72* DNA, and C represents no template control.

### Assessment of ASFV/crRNA-Cas12a complex cleavage specificity

We tested the Cas12a cleavage specificity to ensure that it was catalytically active in the presence of both target-specific crRNA and RPA amplicons. Fluorescent signals were detected only in the presence of ASFV amplicons and ASFV/crRNA-Cas12a complex but not in the presence of off-target crRNA or in the reaction lacking the target amplicon (Fig. 3).

**FIG 3.**
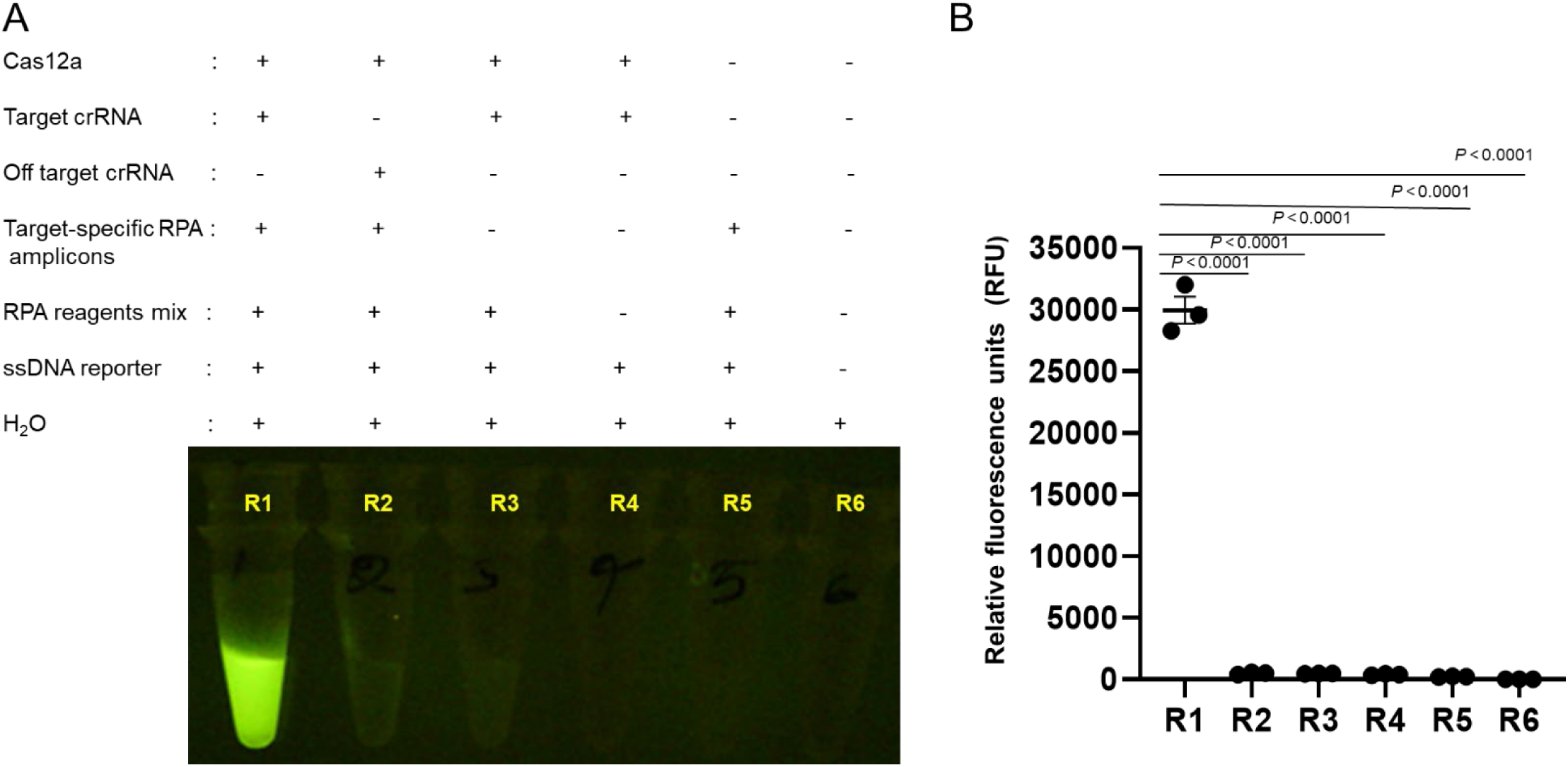
Evaluation of the cleavage specificity of the crRNA-Cas12a complex. A. Schematic representation of the reaction process. A. A visualization of the fluorescence observed under blue light. B. The quantification of the fluorescence signal observed in (A) using a Tecan Spark multimode microplate reader. R1-R6 represent the components present in each tube that are shown in Fig. 3A. The data are mean ± SEM from the three replicates. P values were calculated using a Student’s t-test.

### Optimization of the RPA-CRISPR-Cas12a assay for visual detection of ASFV

To determine the effects of incubation time on the resulting fluorescent signal, the RPA reaction was performed for 2, 5, 10, 15, 20, 25, and 30 minutes. The resulting amplicons were subjected to the CRISPR reaction, and the fluorescence signal was observed incrementally. A clear visible fluorescence signal was observed within 10 minutes consisting of a 5-minute RPA amplification, followed by a 5-minute CRISPR reaction (Figs. 4A and 4B). The 20-minute RPA product showed increased fluorescence with longer CRISPR incubation, which was visualized under blue light. Maximum fluorescence was observed with a 20-minute CRISPR incubation (Figs. 4B and 4C). These results confirm that the optimal reaction time is 40 minutes (including both RPA and CRISPR reactions) to visualize the most intense fluorescence signal.

**FIG 4.**
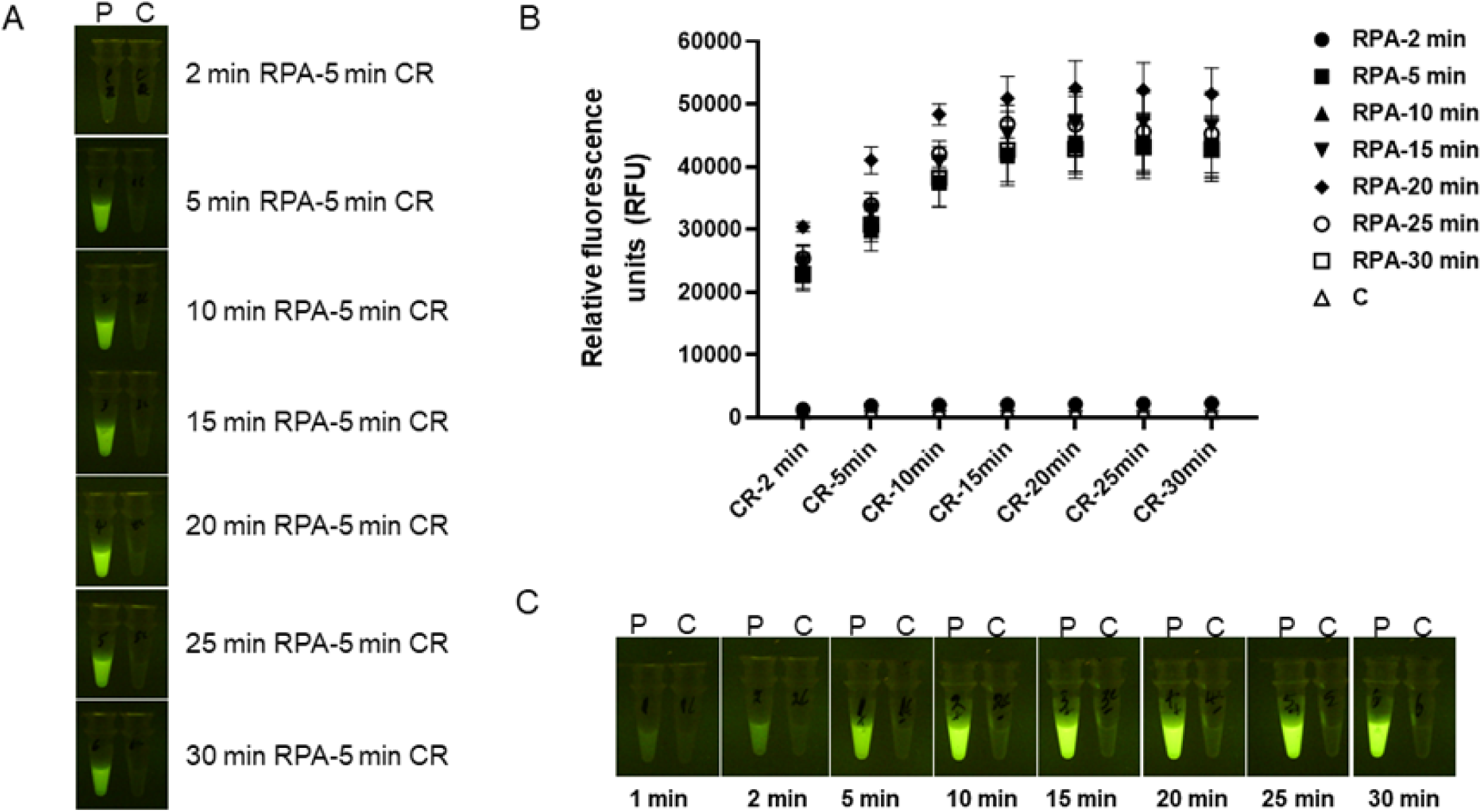
Optimization of RPA and Cas12a reaction time. RPA and CRISPR reactions were assessed with incubation times from 2-30 minutes. A. Resulting fluorescence was assessed with 2-30 minutes RPA reactions and a constant CRISPR reaction of 5 minutes. B. Fluorescence signals were quantified using a Tecan Spark multimode microplate reader. C. The 20-min RPA reaction amplicons were employed in the CRISPR reaction for 1, 2, 5, 10, 15, 20, 25, and 30 minutes, and the resulting florescence was observed by visualization under a blue light transilluminator. The data are mean ± SEM from the three replicates. P represents *p-p72* DNA; C represents no template control; and CR represents the CRISPR reaction.

### Assessment of RPA-CRISPR-Cas12a-assay sensitivity of the ASFV virus target

To determine the analytical sensitivity of the assay, the *p-p72* recombinant ASFV DNA was 10-fold serially diluted with water and tested using the RPA-CRISPR-Cas12a assay and by qPCR. The results showed the limit of detection (LOD) was 10^2^ copies/reaction in both assays (Fig. 5). The 10^2^ copies, equivalent to 7 attograms/uL, exhibited clear visible fluorescence in the RPA-CRISPR reaction. These findings suggest that the RPA-dependent CRISPR-Cas12a-mediated diagnostic tool is comparable to the qPCR assay for detecting ASFV.

**FIG 5.**
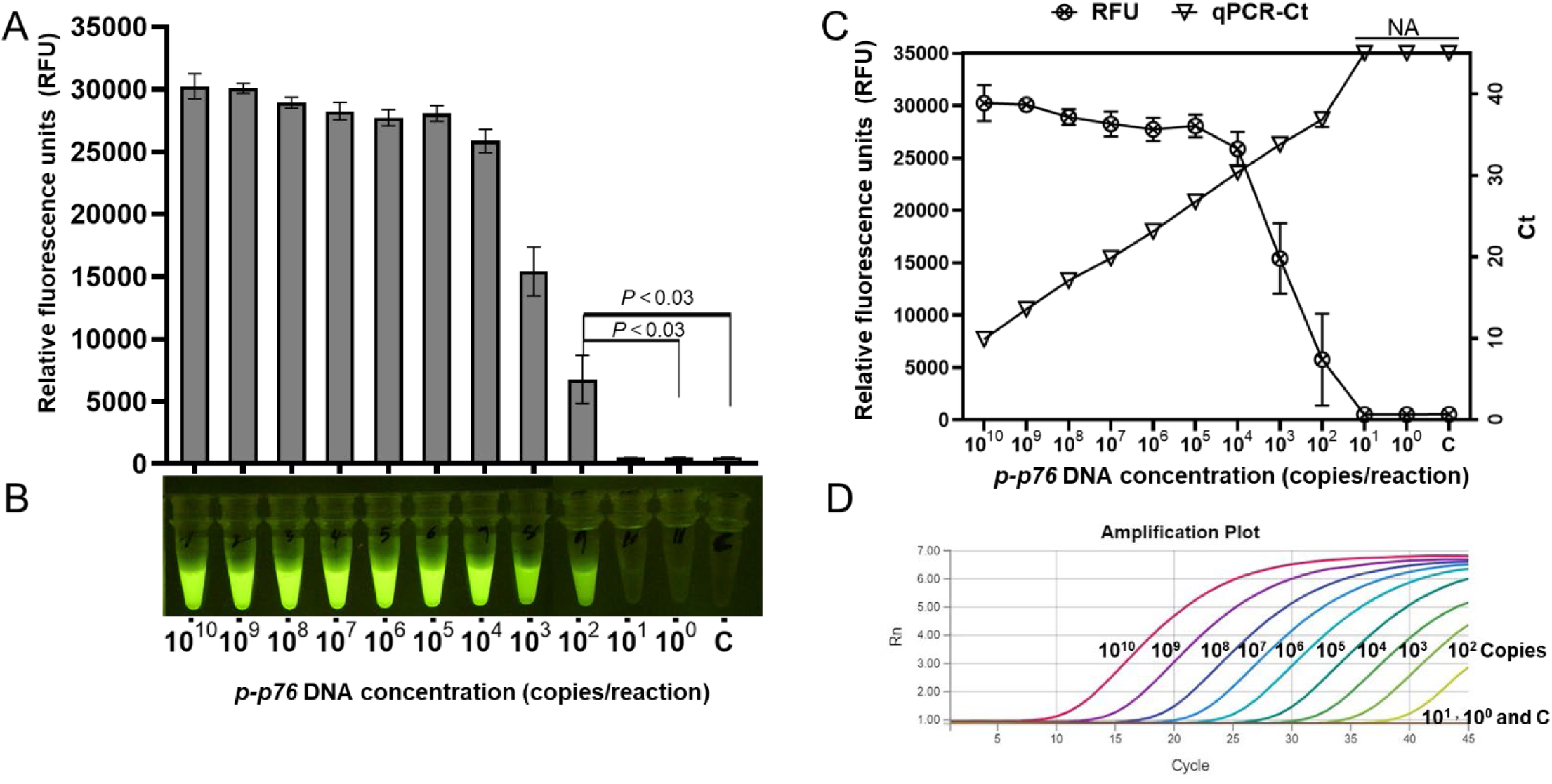
RPA-CRISPR-Cas12a analytical sensitivity to detect ASFV DNA. The sensitivity of the RPA-CRISPR assay was tested using 10-fold serially diluted *p-p72* DNA. A. Fluorescence signals were quantified using a Tecan Spark multimode microplate reader. B. The resulting florescence was observed by visualization under a blue light transilluminator. C. Comparison of the sensitivity of the CRISPR-Cas12a reaction fluorescence with qPCR Ct values. D. The sensitivity of the qPCR assay using 10-fold serially diluted *p-p72* DNA. The data are mean ± SEM from the three replicates. P values were calculated using a Student’s t-test. NA represents no amplification. C represents no template control.

### Assessment of the RPA-CRISPR-Cas12a-assay specificity for the detection of ASFV

To test the analytical specificity of the RPA-CRISPR-Cas12a reaction in detecting the ASFV target region, we used *p-p72* DNA, DNA from SwPV and MVAV, and cDNA from RNA viruses such as PEDV, PHEV, BVDV, and PRRSV. The RPA-CRISPR-Cas12a reaction produced a strong fluorescence signal for the *p-p72* positive control, while no fluorescence signal was observed for the DNA or cDNA of other viruses (Fig. 6). The results indicate that the ASFV-specific RPA-CRISPR reaction did not cross-react with other viral species, successfully identifying the ASFV target by generating a strong, visible green fluorescence signal.

**FIG 6.**
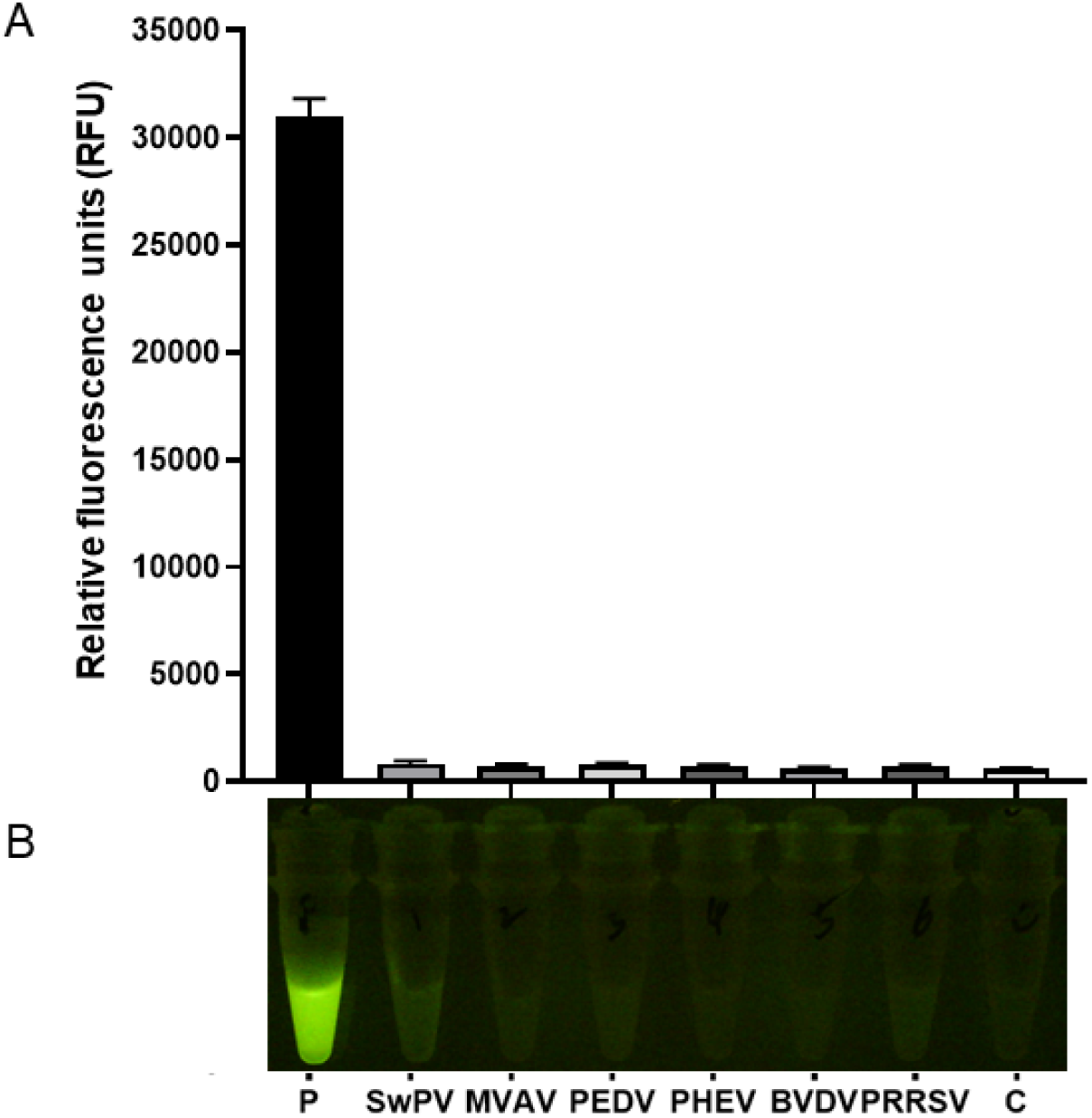
Determination of the RPA-CRISPR-Cas12a analytical specificity. A. Fluorescence signals were quantified using a Tecan Spark multimode microplate reader. B. The resulting florescence was observed by visualization under a blue light transilluminator. The data represent the mean ± SEM of the three replicates. P represents *p-p72* DNA, and C represents no template control.

We mixed *p-p72* DNA into three different sample matrices to assess whether cellular inhibitors from these matrices affect the accuracy of the RPA-CRISPR-Cas12a assay. All spiked DNA samples exhibited a clear, visible fluorescent signal, while the unspiked DNA samples and control samples showed no visible fluorescent signal (Fig. 7A and B). These results correlated with qPCR results (Fig. 7B) indicating that the RPA-CRISPR-Cas12a assay robustly detects ASFV.

**FIG 7.**
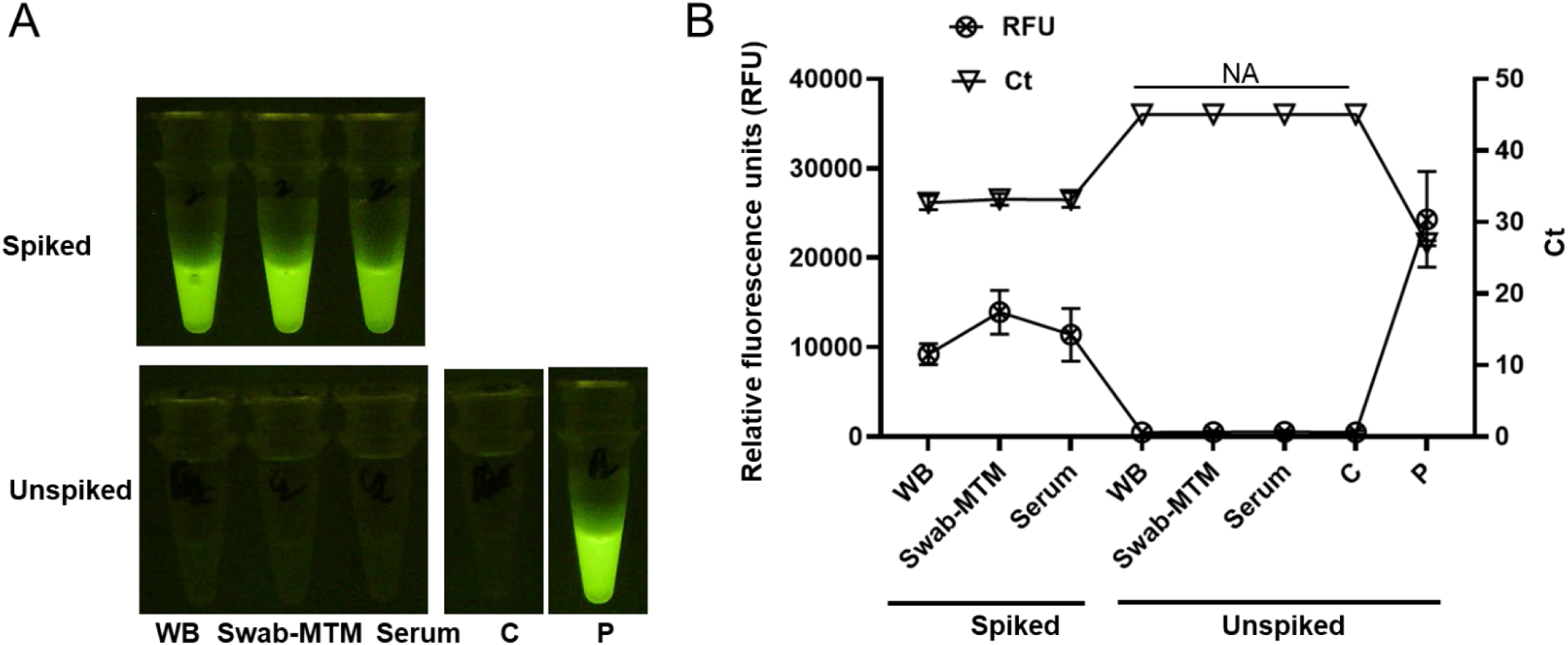
Testing the precision of an RPA-CRISPR-Cas12a assay using *p-p72* DNA spiked sample matrices. The *p-p72* DNA was spiked in different sample matrices and tested with the RPA-CRISPR-Cas12a assay. A. The resulting florescence was observed by visualization under a blue light transilluminator. B. Fluorescence signals were quantified using a Tecan Spark multimode microplate reader. The data represent the mean ± SEM of the three replicates. C represents no template control, P represents *p-p72* DNA, which was not spiked as a positive control, NA represents no amplification, and WB represents whole blood.

### Diagnostic sensitivity and specificity of the RPA-CRISPR-Cas12a assay to detect ASFV samples

To assess the reliability of the RPA-CRISPR-Cas12a assay for visual detection of ASFV, we screened 66 suspect blood samples from the suspect farms and 39 negative surveillance blood swab-MTM samples. Of the 66 suspect blood samples, 46 showed intense fluorescence, while one sample had a notably less intense resulting fluorescence (Table 2, Supplementary Figure 1, Supplementary Table 1). Similarly, qPCR testing identified 47 positive samples; however, the single sample with the lower fluorescence in the RPA-CRISPR test had a Ct value of 37 (Supplementary Table 1). The remaining 19 suspect blood samples had no visible fluorescence in the RPA-CRISPR-Cas12a assay and were negative by PCR (Table 2, Supplementary Figure 1, Supplementary Table 1). Additionally, the 39 negative surveillance blood swab-MTM samples showed no visible fluorescence (Table 2, Supplementary Table 2). The results of the RPA-CRISPR-Cas12a assay with both suspect and surveillance diagnostic samples were consistent with qPCR results. The calculated confidence intervals for diagnostic sensitivity and specificity were 98.3% and 100%, respectively (Table 2).

**TABLE 2.**
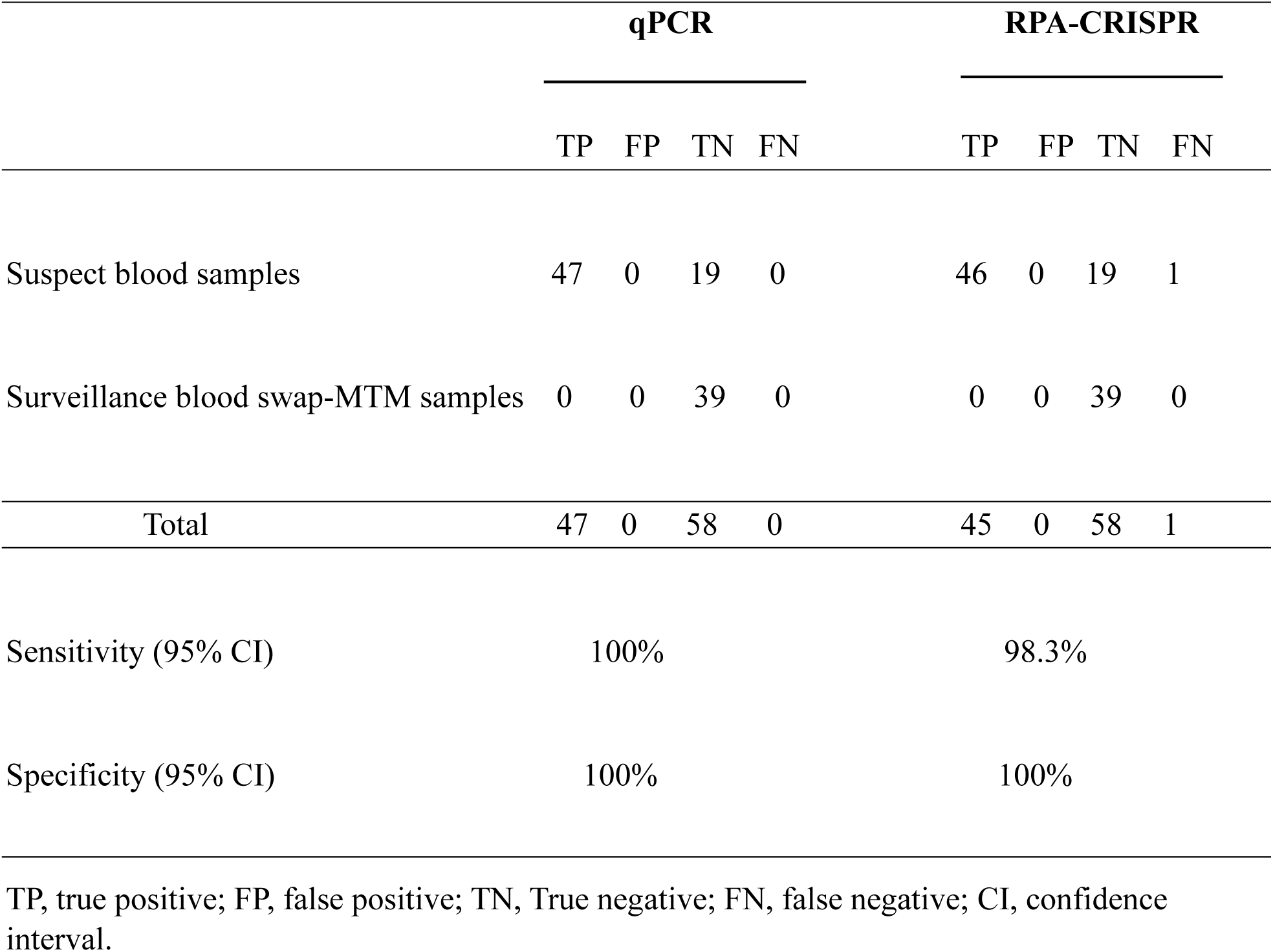
The sensitivity and specificity compare between qPCR and RAP-CRISPR-Cas12a diagnostic assay results from diagnostic samples.

---

**Supplementary FIG 1.**
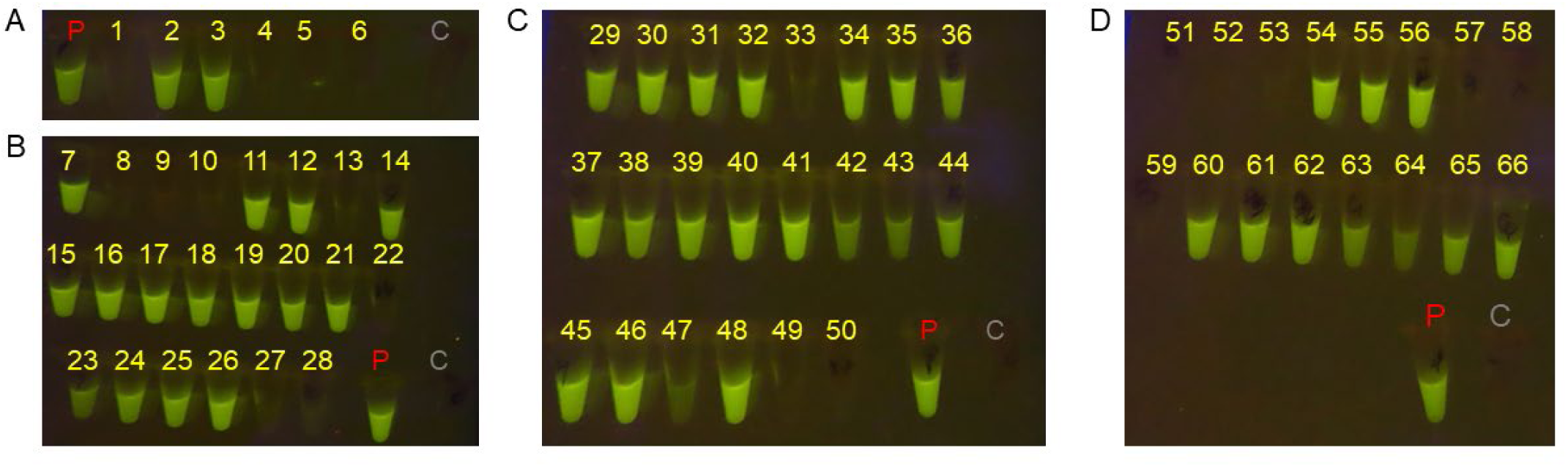
The suspect blood samples were tested for ASFV detection. A-D. The RPA-CRISPR-Cas12a assay resulted in green fluorescence visualized under a blue light transilluminator, and images were taken using a digital camera. 1 to 66: the number of suspected diagnostic blood samples that were tested. P represents *p-p72* DNA used as a positive control, and C represents no template control.

## DICUSSION

ASF is a highly contagious disease of swine that leads to economic losses and disrupts trade. The recent ASFV outbreak in the Hispaniola threatens the US, the third largest global pork producer, due to its proximity and role in North American trade routes. Currently, US mitigation strategies focus on preventative measures such as biosecurity as there is no licensed and effective ASFV vaccine available (Borca et al., 2024; Bohorquez et al., 2023). The standard laboratory molecular diagnostic assay to detect and survey for ASFV is qPCR; however, qPCR is not suitable for field-use as it requires costly equipment, infrastructure, and training to perform successfully.

Isothermal-based tests like RPA and LAMP have the potential to overcome these challenges but suffer from high false positive rates in field evaluations (Miao et al., 2019; Zhai et al., 2020). Adding CRISPR to this isothermal platform may address these issues due to its ability to specifically identify target nucleic acids. Previous studies coupling RPA and CRISPR-Cas12a for ASFV detection have shown promise, improving the false positive rates of the isothermal platforms; however, no study has yet optimized the assay or improved its sensitivity to a level suitable for field surveillance.

In this study, we developed an RPA-CRISPR-Cas12a-mediated *p72* ASFV visual detection assay based on the sequence utilized for the qPCR assay. We optimized and validated the assay using field diagnostic samples for visual detection. Selection of target-specific crRNA sequences is crucial to create efficient and specific crRNA-Cas-mediated cleavage activity (Konstantakos et al., 2022; Corsi et al., 2022). We assessed two crRNAs targeting *p72* of ASFV, one of which successfully showed increased fluorescence when detecting the ASFV target to a higher degree than previously published assays both in analytical and diagnostic validation of sensitivity and specificity.

These results confirm the importance of target-specific crRNA sequence selection to the developed CRISPR assay’s sensitivity and specificity thresholds. We also confirm that neither the Cas12a alone nor the crRNA-Cas12a complex is catalytically active enough to cleave the ssDNA reporter. Our study assessed a single-tube, two-step RPA-CRISPR-Cas12a assay resulting in higher sensitivity and specificity than the previously published single-step approaches for ASFV detection (Qin et al., 2022). We followed this approach to avoid cross-contamination, reduce false positives, and maximize the visual detection limit for ASFV in diagnostic samples.

This approach enabled visual detection in as little as 7 minutes (5 minutes RPA reaction and 2 minutes CRISPR-Cas12a reaction) with peak visual fluorescence observed at 40 minutes (20 minutes RPA reaction and 20 minutes CRISPR-Ca12a reaction). The previous study showed clear visual fluorescence in 40-60 minutes to detect ASFV using positive control DNA but had lower sensitivity and lacked validation of its reliability (Qin et al., 2022). We used 40-minute reaction time as it was the time in which peak fluorescence was visualized in the assay to ensure low-titer virus was detected in the results. This 40-minute reaction time was much quicker than qPCR in the detection of ASFV (Zsak et al., 2005).

Previous studies using isothermal and CRISPR assays have shown variable success in meeting the standards set by the gold-standard ASFV qPCR assay, including the limit of detection (LOD) of 10^2^ genome copies per reaction, and have not been tested on diagnostic samples (Fu et al., 2021; Qin et al., 2022). The sensitivity limit can vary depending on the type of template (recombinant plasmid DNA, gDNA, or cDNA), the choice of target region, the efficiency of the crRNA and Cas enzymes, and the detection methods used (visualization, machine-reading fluorescence signals, or lateral flow) (Tian and Zhou, 2023). Previous studies reported LODs between 1 and 10^4^ copies/uL when using an isothermal reaction coupled with CRISPR (Ren et al., 2021; Fu et al., 2021; Qin et al., 2022; Tian and Zhou, 2023).

Our sensitivity analysis showed that the RPA-CRISPR assay could detect 10 copies/uL, both visually and quantitatively, comparable to the qPCR LOD. In contrast, a similar, recently published RPA-CRISPR assay reported a visual LOD of 10^4^ copies/uL (Qin et al., 2022). Our results not only match qPCR sensitivity, but the assay is highly specific, as other viral DNA or RNA did not cross-react in our assay. Furthermore, we tested various matrices (whole blood, serum, or blood swab in MTM) and found they did not interfere with the RPA-CRISPR-Cas12a assay.

The assay developed herein is both sensitive and specific despite the variable sample matrices. It meets the standards set by the World Health Organisation’s (WHO) ASSURED criteria (affordable, sensitive, specific, user-friendly, rapid, equipment-free, and delivered) to ensure equal access across the globe (Ghouneimy et al., 2023). The RPA-CRISPR-Cas12a ASFV assay does not require expensive equipment, sophisticated facilities, or specialized training, can be run in as little as 7 minutes, and meets the guidelines set out by WOAH (terrestrial manual, chapter 1.1.6, 2024) for diagnostic tests.

In this study, we followed WOAH guidelines to validate diagnostic sensitivity and specificity using both true positive and negative samples (WOAH, terrestrial manual, chapters 1.1.6 and 3.9.1, 2024). To our knowledge, this is the first study to test an appropriate number of diagnostic samples, providing consistent results with qPCR standards for RPA-CRISPR-Cas12a ASFV diagnostic testing. Overall, the results of these experiments support the use of a rapid, affordable, point-of-care ASFV test, potentially during an outbreak in the US.

## ACKNOWLEDGEMENTS

The authors thank Alethea Fry and Dr. Alexandra Scupham from the USDA Center for Veterinary Biologics for providing various viral DNA and cDNA samples. We are grateful to Dr. Gleeson Murphy for his kindly critical review and for his useful suggestions. We thank Dr. Elizabeth Lautner and Dr. Ping Wu for reviewing the manuscript and their useful comments. We also thank the USDA-APHIS Wildlife Services for supplying negative diagnostic samples as part of the ASFV feral swine surveillance program. Additionally, we appreciate the Laboratorio Veterinario Central (LAVECEN) in the Dominican Republic for providing ASFV suspect field samples. The findings and conclusions in this publication are those of the authors and should not be construed to represent any official USDA or U.S. Government determination or policy.

## AUTHOR CONTRIBUTIONS

SK, RP: Conceptualization and writing; SK: Methodology and analyzed the data; JT: Methodology; SRA: Feedback, review and editing, funding acquisition; KS: Feedback, review, editing and funding acquisition; RP: Supervision. All authors reviewed, commented on and approved the manuscript.

## FUNDING

This project was funded by the USDA Animal and Plant Health Inspection Service through the National Veterinary Services Laboratories

## CONFLICT OF INTEREST

The authors declare no conflict of interest.

## Notes

### Competing Interest Statement

The authors have declared no competing interest.

